# platpy: A spatial-first framework for multi-layer spatial transcriptomic analysis

**DOI:** 10.64898/2026.05.26.727917

**Authors:** Thomas M Maynard

**Affiliations:** Fralin Biomedical Research Institute at Virginia Tech Carilion 2 Riverside Circle, Roanoke, Virginia, 24016, USA

**Keywords:** spatial transcriptomics, Visium HD, Xenium, Python, spatial analysis, bioinformatics

## Abstract

**Background:** The emergence of accessible spatial transcriptomic platforms such as 10x Genomics Visium HD and Xenium has created demand for analysis tools that can handle the complexity and scale of spatial datasets. Current frameworks approach spatial data primarily as an extension of single-cell RNA-seq pipelines, where spatial coordinates are retained as metadata rather than treated as a first-class organizing principle. As a result, common tasks such as multi-modal data alignment, region-of-interest selection, and cross-resolution visualization require manually managing disparate data types, coordinates, and scales, making spatial analysis unnecessarily time-consuming and error-prone.

**Results:** We present platpy (<u>p</u>ipeline for layered <u>a</u>nalysis of transcriptomics), a Python-based “spatial-first” framework that treats absolute physical micron coordinates as the organizing principle for all data types. All data – morphology images, transcript point clouds, expression matrices, segmented cells, and user-defined regions – are stored as typed objects (“Channels”) that carry their own spatial metadata, keeping all layers in automatic registration regardless of platform, resolution, or analysis operation. Two complementary interfaces simplify access to underlying data: the *ViewPort*, a compositing engine for efficient multi-channel visualization, and the *DataPort*, which extracts raw data in its native format for downstream analysis. A set of spatial analysis tools demonstrates the practical benefits of the framework, including ROI-based expression binning, cortical unfolding, and sub-micron fine alignment of transcript and image data. The use of modern Python data management methods helps maintain the efficiency of the framework, allowing for quick visualizations and analysis with a low memory footprint.

**Conclusions:** Platpy is designed to complement rather than replace widely used tools in the spatial analysis ecosystem (scanpy, squidpy, CellPose, StarDist), by handling the spatial mechanics of large datasets so that the analyst can focus on the biology. Platpy is freely available under the MIT license at https://github.com/maynardt/platpy.

## Background

The emergence of accessible spatial transcriptomic platforms such as 10X Visium HD [1] and Xenium [2] has caught the attention of biologists across disparate fields. The merging of high-resolution imaging and single-cell level transcriptomic data promises the power of molecular genetics to investigate cellular function, while providing spatial insights that fuel cell biological investigation. However, the complexity and size of spatial datasets have made spatial analysis time consuming, resource intensive, and unnecessarily difficult. Simple questions can require hours of tricky coding to align data layers of different modalities, and processing multi-gigabyte datasets can bog down high-end workstations for hours.

Current analysis frameworks approach spatial data primarily as an extension of single-cell RNA-seq pipelines: transcripts are aggregated into cell-by-gene count matrices, with spatial coordinates retained as metadata. However, the handling of the spatial aspects of analysis is relatively cumbersome. Published workflows using scRNA-seq based tools such as Scanpy [3] or Seurat [4, 5] require that the user manually align transcript and image layers, and manually manage different coordinate systems and array dimensions, making simple tasks like selecting an ROI a surprisingly challenging and error prone task.

To simplify spatial analysis, we have developed platpy (<u>p</u>ipeline for layered <u>a</u>nalysis of transcriptomics), a Python-based “spatial-first” framework that treats physical micron coordinates – rather than array indices – as the organizing principle for all data types. Platpy is not designed as a replacement for widely used tools in the spatial ecosystem (Scanpy, Squidpy [6], CellPose [7], StarDist [8], etc.); rather, it is designed as a scientist-friendly layer that cooperates with these tools, handling the mechanics of working with large spatial datasets so that the analyst can focus on the biology.

## Implementation

Platpy is built around a model in which all data types for an experimental sample are organized within a single container, the *SampleSet*. A *SampleSet* holds a collection of data sets, each of which is stored as an object called a *Channel*. Four types of channels are currently defined: *ImageChannel* (morphology and immunofluorescence images), *OmicChannel* (transcript point clouds or spot-level expression matrices), *CellChannel* (segmented cell data), and *ROIChannel* (user-defined spatial regions and annotations). Importantly, each channel object carries with it the necessary information for mapping to physical micron coordinates of the source material. This mapping is done via a *SpatialMetadata* object that encodes the data’s position and scale as an affine transformation matrix. This matrix is the single source of truth for where data lives in physical space: all pixel-to-micron and micron-to-pixel conversions flow through it, ensuring that channels of different types and resolutions remain in perfect registration. Multiple channels of each type can co-exist; for example, a single sample might carry several *ImageChannels* representing different immunofluorescence markers, multiple *ROIChannels* defining independent regions for different analyses. In principle separate *OmicChannels* could also carry parallel proteomic data or ATAC-seq data generated from the same tissue for integration at the cell/tissue level. Platpy currently supports 10x Genomics Visium HD and Xenium platforms, but the channel architecture is designed to be extensible to additional spatial platforms without changes to the analysis layer.

A major challenge in spatial analysis is the integration of multiple layered data types, each stored at different scales and resolutions, and accessed in different ways. To allow analysis to focus on the data, rather than mechanics of handling disparate data types, platpy introduces the *ViewPort*, which is a simplified compositing engine for visualizing spatial data. Users define a view by specifying a center point and field of view in absolute microns, add any combination of channels as layers, and then add optional orientation parameters or display options (rotation, transparency, color maps, etc.). Rendering is lazy – layers are composited only when explicitly requested, at whatever resolution the user specifies. The resulting render is itself a registered *ImageChannel*, meaning it retains its micron-space coordinates and can be passed directly to downstream analysis. A key use case is interactive ROI selection: a screen-resolution render can be handed off to Napari [9] for visualization and user interaction. Any ROIs drawn in pixel space are automatically translated back to absolute micron coordinates by the *ViewPort*, feeding directly into data extraction or spatial analysis without manual coordinate management.

While the *ViewPort* is designed for visualization, its analytical counterpart, the *DataPort*, handles extraction of raw data from the same spatial frame. Where a *ViewPort* renders a screen-resolution composite, a *DataPort* extracts data in its native format – images are returned as unmodified arrays, and transcriptomic data remains in native form (whether Visium HD bins, Xenium point-clouds, or raw 16-bit image arrays) to prevent the introduction of resampling or normalization artifacts before analysis. A *DataPort* is initialized directly from micron coordinates (for example, directly from a *ViewPort*, or from an *ROIChannel* selecting a structure for analysis), guaranteeing that the visual selection and the data extraction operate on the same ROI. The *DataPort* prepares data for analysis, but it is not an analytical tool itself. Analysis functions such as cell segmentation (CellPose, StarDist) are passed into the *DataPort* rather than implemented by it: the *DataPort* handles the spatial bookkeeping, and the science stays in the calling code. Results are returned with their absolute micron-space coordinates attached, so segmentation masks and other outputs can be mapped to any other data layer seamlessly. For large ROIs that exceed available memory, *DataPort* handles automatic tiling, with each tile carrying its own spatial extent so that results can be reassembled in physical space without additional coordinate management.

To illustrate the practical benefits of the unified micron-space framework, we have included a set of spatial analysis tools (Figure 2) that demonstrate the relative simplicity of spatial approaches within platpy, freeing the scientist from managing different data types and resolutions. First, a set of three ROI-generating tools are provided that can be used to quickly generate histogram bins from different tissue structures: Axis binning, to slice a structure into equal-width bins (e.g. to illustrate dorsal-ventral gradients); Contour-shell binning, to slice a structure into “onion-skin” layers (e.g. to demonstrate “core” vs “periphery” gradients); and a “depth bin” approach (designed for illustrating layer-based gradients in the cerebral cortex or similar layered structures). While such binning approaches are technically possible in standard pipelines, these demonstrations show that the platpy framework simplifies the implementation, and the user can generate useful analysis in a few lines of code without concern for manual coordinate management or data handling. An additional tool implements a “cortical unfolding” algorithm that re-maps layered cortical cell expression data from a curved segment of cortex to a uniform rectangular slice. Finally, we demonstrate a workflow that implements a “fine alignment” function that generates a sub-micron level correction to precisely map Visium HD spot data to the paired immunofluorescent images, allowing for accurate cell segmentation across data types. These tools are not intended to be exhaustive; rather, they demonstrate that once all data types share a common physical coordinate system and unified methods of accessing the underlying data, spatial analyses that are cumbersome in array-index-based frameworks become straightforward to implement.

**Figure 1.**
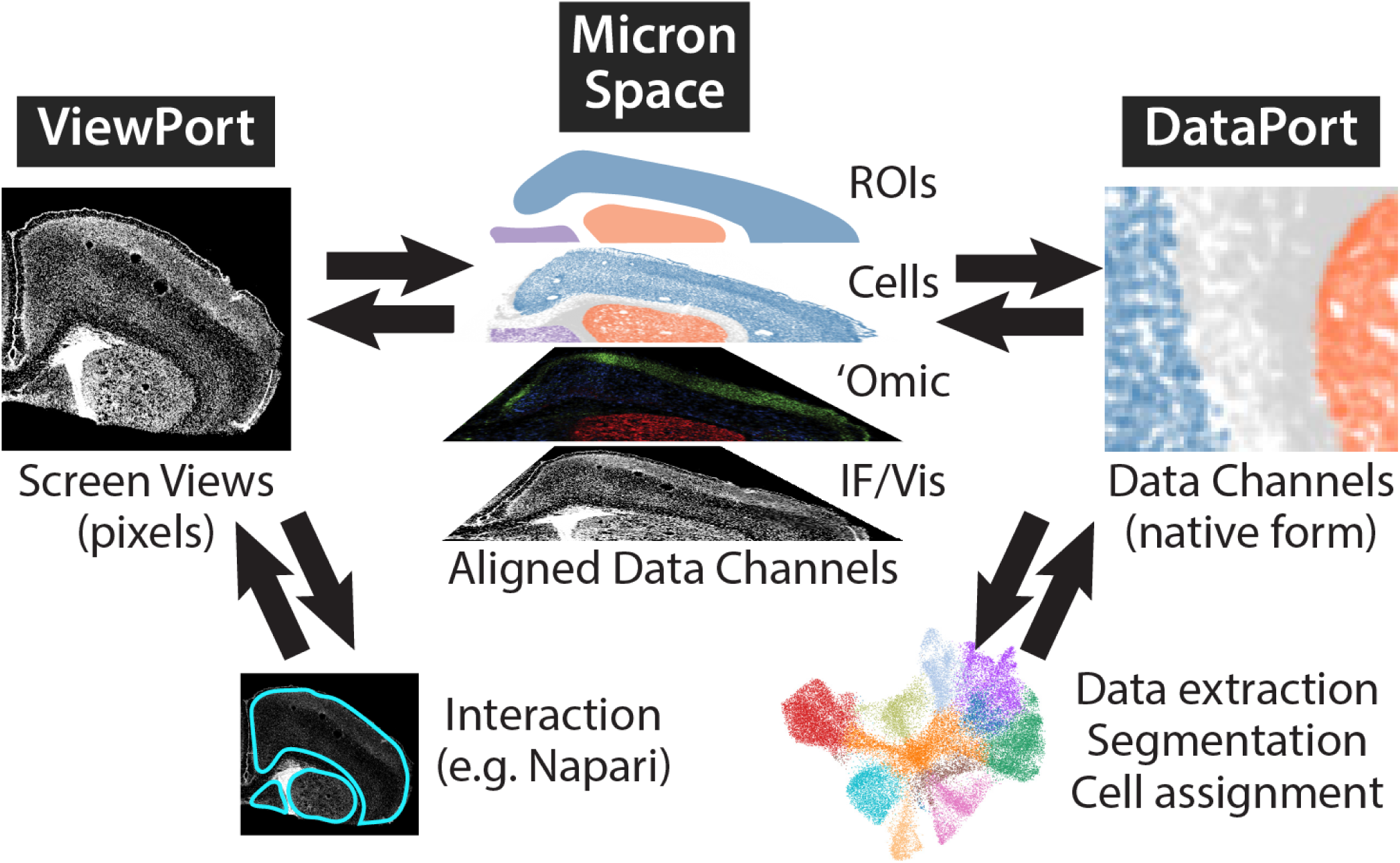
Overview of platpy architecture and data handling. Spatial data in platpy is defined by its location in micron space, referring to its spatial coordinates on the original sample tissue. Data objects (image arrays, transcriptomic data in spots/bins or point-cloud form, or derived data like segmented cells) as well as annotations like ROIs are placed in self-contained objects (“Channels”) that contain spatial metadata that map them to micron space. Two primary interfaces exist for viewing and working with layered data: the *ViewPort*, which provides screen-resolution renderings of images and data, and the *DataPort*, which provides access to raw data for analysis. Importantly, interactions with the *ViewPort* and *DataPort* are bi-directional: A screen-resolution view of an ROI can be used interactively (e.g. by selecting ROIs in Napari); the results are passed back through the *ViewPort* to translate the same ROI to absolute micron coordinates. Likewise, the *DataPort* can be used to process raw data; and results are passed back through the *DataPort* to transform spatial results back to micron space.

**Figure 2.**
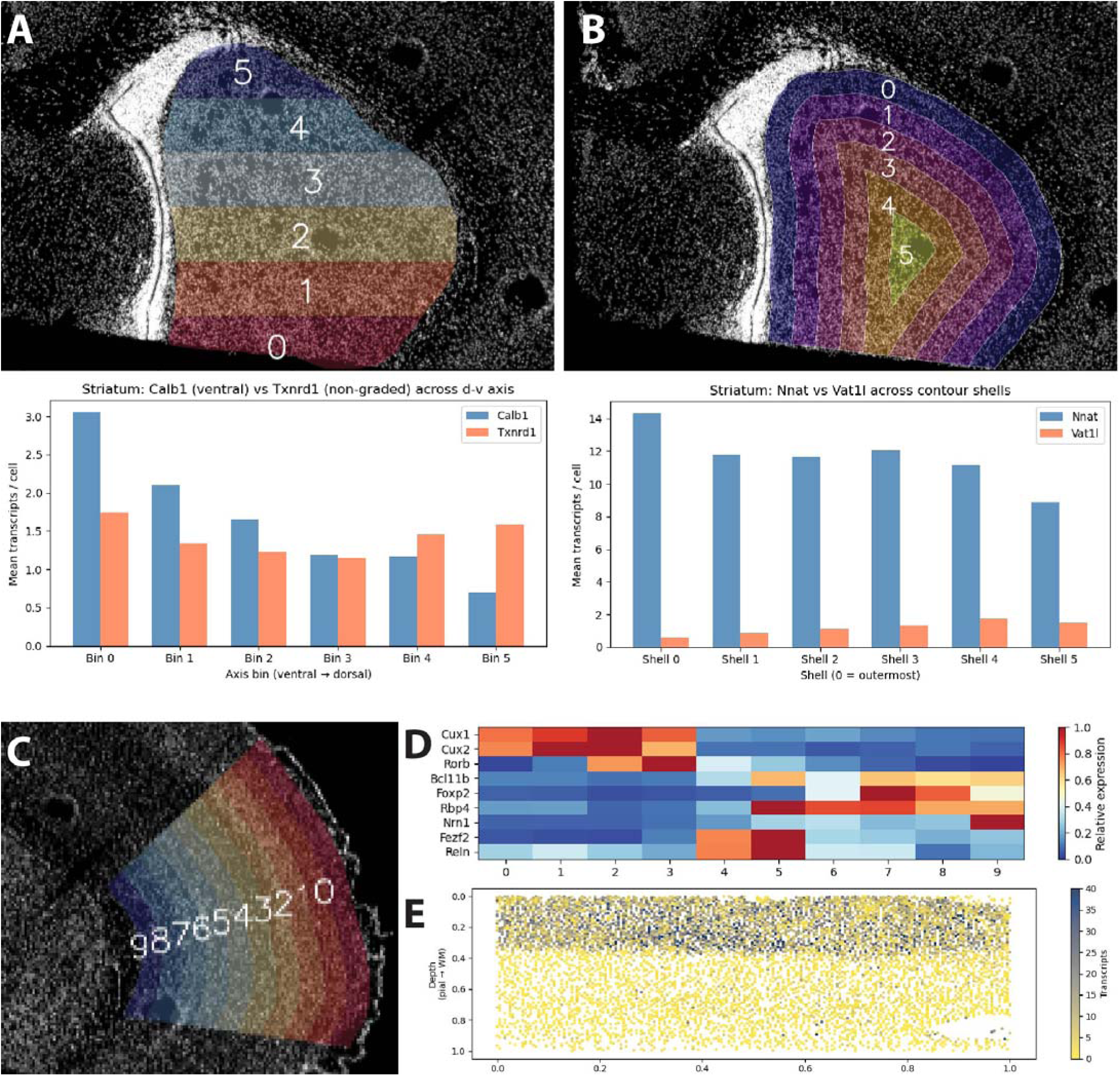
Examples of spatial analysis tools demonstrated in provided vignettes. A. Generation of fixed-width ROI bins from a sample ROI (P5 striatum) to generate a histogram displaying dorsal-ventral expression gradients. B. Generation of contour-aware “onion skin” ROI bins to demonstrate core-periphery gradients. C. Contour-aware mapping of “depth bins” on a section of cerebral cortex. D: Heat map of cortical bins from C; note high level expression of upper-layer specific markers (Cux1, Cux2, Rorb) mapping to superficial bins; remaining markers are expressed in subsets of lower-layer neurons. E. Remapping of segmented cells from same section of cortex as C to produce an “unfolded cortex” map to illustrate cortical layering; color map represents combined expression of upper-layer markers Cux1 and Cux2. All analysis and visualizations are aided by platpy’s workflow that maps data layers to a uniform coordinate space.

**Figure 3.**
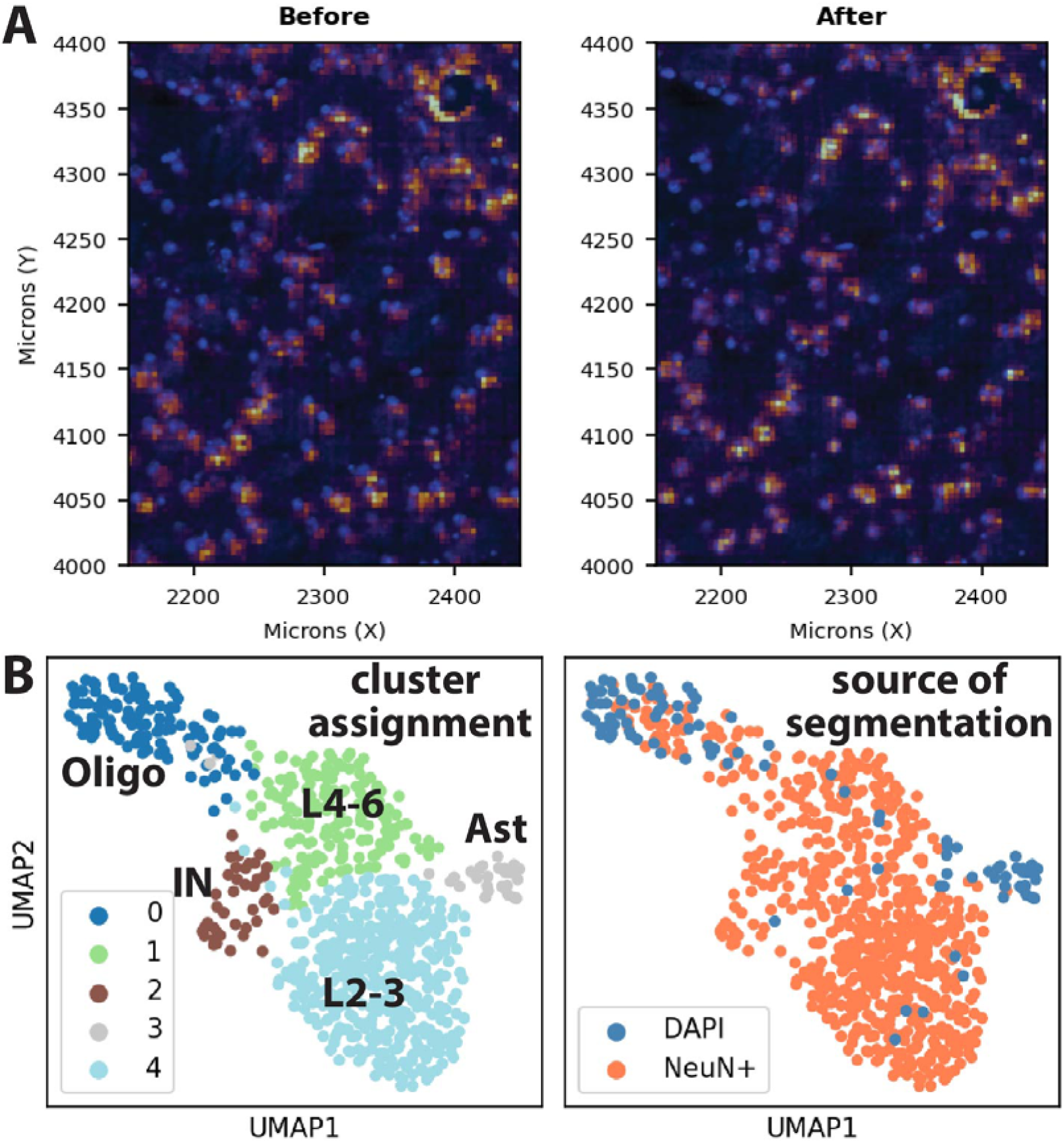
Cell-level alignment of Visium HD spot data facilitates segmentation via immunofluorescence channels. A. Before-and-after view of Visium HD data (reds) vs. DAPI image (blue); note that Space Ranger provides coarse alignment, but a cell’s transcripts may only partially overlap its nuclear image. After alignment, a correction factor is applied to spot-level data to provide clean alignment. B. Assignment of identities to segmented cells: Using the aligned data, two rounds of segmentation were performed: first, neurons were assigned based on NeuN immunofluorescence [12]; assigned regions were then masked and then residual cells were assigned by segmentation to DAPI labeling. Two UMAPs are shown, the first illustrating the key identified cell types, and the second displaying which cells were identified as NeuN+ vs. DAPI. The DAPI-only profiles are mostly confined to non-neuronal (glial) cell types as expected. Some alignment error is likely due to overlap of cell nuclei, particularly oligodendrocytes which have small nuclei and are closely associated with neuronal processes. Abbreviations: Ast: astrocytes, Oligo: oligodendrocytes, IN: cortical interneurons, L2-3: upper-layer cortical pyramidal neurons (e.g. Cux1/2+), L4-6: lower-layer cortical pyramidal neurons.

Platpy is designed to operate efficiently on standard research hardware. Lazy loading via Dask [10] and Zarr [11] ensures that image and transcript data are read from disk only when needed, keeping the memory footprint low during analysis. To speed image visualization, multi-resolution image pyramids are generated automatically, and the *ViewPort* selects the appropriate resolution at render time. Furthermore, the affine matrix-based method for translating image and data arrays to micron-space and back is exceptionally efficient (and highly optimized by modern CPUs/GPUs), allowing almost instantaneous screen views of images regardless of scaling, rotating, or coordinate flips. Together, these optimizations allow the full analysis and visualization workflow to run on a standard laptop; all vignettes accompanying this paper were tested and run efficiently on a 16GB Apple MacBook Air (M2), except for initial data loading which requires proportionally more memory for large datasets to generate optimized Dask/Zarr files. Full-scale analysis of multiple samples will likely require significantly more resources; nevertheless, the efficiencies provided by platpy’s core functions will help facilitate performing full-scale analyses on standard laboratory-grade workstations (e.g. 32-64G RAM) without the need to resort to extremely high specification hardware or high-performance compute clusters.

## Discussion

The key design goal of platpy was to generate spatial tools that seamlessly integrate with the established Python single-cell and spatial analysis ecosystem. Rather than reimplementing functionality that already exists in mature, well-supported, and widely-used tools, platpy is designed to sit alongside them and handle the spatial mechanics of data access and visualization. The goal of the “spatial-first” framework is to facilitate spatial tasks: visualization, ROI selection, and spatial data handling, without manual coordinate translation or format conversion. This design philosophy reflects a broader observation: the barrier to spatial transcriptomic analysis is often not the analysis itself, but the overhead of managing large, multi-modal datasets before the analysis can even begin. By addressing this overhead directly, platpy is intended to make spatial transcriptomics more accessible to biologists across fields – including those without a dedicated bioinformatics infrastructure – and to reduce the time between data acquisition and biological insight.

In its current form as an initial release, there are several areas of the framework targeted for future improvement. While image handling is highly optimized in this version, additional optimizations are still planned for handling large transcriptomic datasets, including spatially “chunking” and indexing spatial data to increase speed and efficiency when examining small ROIs. The pipeline for loading raw 10x data is particularly inefficient; however, this is not a focus of development: it is assumed that loading of raw data can be a “one and done” part of an analysis project, so optimization is best focused on more extensively used functions. Longer term, the channel architecture is designed to accommodate additional spatial platforms as they emerge — the rapidly evolving spatial transcriptomics landscape makes this an important design consideration for any framework intended to remain useful over time.

## Conclusion

Platpy is a spatial-first utility for working with large spatial datasets. It is designed to reduce the barriers to multi-modal spatial analysis by providing a simple, intuitive interface to the scientist by abstracting away coordinate conversion and handling of multi-gigabyte data files. It is intended to be integrated within the well-established set of Python tools for spatial analysis (Scanpy, Squidpy, etc.), and to complement these tools rather than replace them. platpy is platform agnostic and has been tested on macOS, Windows, and Linux. It is freely available under the MIT license at https://github.com/maynardt/platpy.

## Acknowledgements

Thanks to Anthony LaMantia for his support in the development of this package, and for assisting with manuscript preparation. The sample Xenium data was generated as part of a spatial transcriptomic analysis that has been submitted for review elsewhere; key authors on this work also include Shah Rukh, Zach Erwin, Daniel Meechan, and Anthony LaMantia.

## Funding

This work was supported by NIH 5R01-HD042182, and by The Red Gates Foundation.

## Availability of data and materials

Code and example vignettes are freely available on GitHub under MIT license at (https://github.com/maynardt/platpy). Sample data sets for both the Visium HD and Xenium examples in this manuscript and in the vignettes are available from Zenodo (doi:10.5281/zenodo.20320691). Xenium raw data is deposited in the GEO database under GSE342913.

## Notes

https://github.com/maynardt/platpy

https://zenodo.org/records/22149474

### Competing Interest Statement

The authors have declared no competing interest.

### Summary of Updates

The package has been renamed in response to resolve a conflict with another unrelated python package.

